# Policy-Enforced Genomic Repositories and Institutional Data Governance for African Pathogen Surveillance with UbuntuBio

**DOI:** 10.64898/2026.11.16.767022

**Authors:** Addo Stephen Tetteh, Richard Owusu Ansah, Kwame Boamah Buabeng, Kwasi Agyenkwa-Mawuli, Daniel Baah, Kofi Sarpong, Anjali Rathod, Juliet Naa Otua Hodgson, Rachel Ama Adadziwa Akorful, Baakwa De-graft Miah, Adwoa Ayewa Ayimadu, E. K. Amoako, Samuel Mawuli Kumordzi, Arnold Abakah, Nana Yaw Tawiah-Yingar, Doreen Akakpo, Rexford Adade, Redeemer Salami Okekale, Elijah Salami Okekale, Marious T. Senyo

## Abstract

Research centers curating pathogen sequence cohorts often operate under constrained bandwidth, multi-institutional data-sharing agreements, and strict biosafety requirements. Global public archives keep released sequences permanently open, external viral portals enforce terms outside local institutional custody, and standard Git Large File Storage (LFS) duplicates identical binaries across repositories without validating taxonomy or geography.

We present **UbuntuBio** (`v1.0.0`), a Git-native, policy-enforced genomic repository platform deployed at https://ubuntubio.org and developed at the West African Centre for Cell Biology of Infectious Pathogens (WACCBIP), University of Ghana. UbuntuBio integrates four mechanisms under a single storage root: (1) bare Git repositories storing dataset cards, sample sheets, and approximately 130-byte LFS pointers backed by prefix-sharded SHA-256 content-addressed storage (CAS) with visibility-scoped wire deduplication; (2) a four-phase commit pipeline recoverable via cross-plane reconciliation that enforces 55 African Union country codes against offline Natural Earth polygons, core MIxS-derived fields, Taxon `9606` rejection, and server-keyed targeted human-marker screening identically across HTTP commits and Git `pre-receive` hooks; (3) institutional custody controls combining Data Use Agreements (DUAs), revocable 900-second RS256 capability tokens, key rotation, tamper-evident hash-chained logs, and externally anchorable RS256 audit checkpoints; and (4) an embedded SQLite WAL and FTS5 catalog filtered by access tier. In single-node evaluation, UbuntuBio classified 18/18 policy boundary fixtures consistently across Python, Git, and HTTP paths; showed 0/265,157 false cross-matches on 21/22 RefSeq pathogen sequences (while cross-matching *Candida albicans* 18S rDNA at 0.113% and *Chlorocebus sabaeus* Alu repeats in ENA stool reads, confirming targeted marker detection rather than verified human DNA decontamination); matched conditional uniform (*n* = 2, 000; 200-seed 0.01% convergence 19.0% vs. 18.13% expected) and interior (*n* = 1, 000) detection theory while detecting all spike-ins at ≥ 0.5% (30/30); halved two-repository disk usage on 613.83 MiB Nigerian *E. coli* runs; passed 27/27 structured-filter correctness checks over 10,000 biosamples and reached graded NDCG@10 = 0.920 on independent free-text BM25 queries with zero private leaks; and recovered cleanly across seven injected phase-transition faults.

## 1 Introduction

Pathogen genomic surveillance expanded rapidly after 2020 across African research networks [1–3]. Yet repository software for curating, validating, and governing raw sequence files at participating laboratories has lagged behind sequencing throughput. Laboratories face two practical bottlenecks. First, large FASTQ and BAM archives waste scarce bandwidth and storage when partner consortia re-upload identical runs across separate namespaces. Second, sample sheets cross institutional boundaries without automated validation for geographic accuracy, metadata completeness, or targeted human DNA markers.

Existing repository platforms impose trade-offs. International archives such as the European Nucleotide Archive (ENA) [4] provide permanent FAIR accessions [5], yet public release ends local custodian control over reuse. Controlled human repositories such as the European Genome-phenome Archive (EGA) [6] run Data Access Committees, but assume cluster infrastructure built for human cohorts. Viral portals such as GISAID [7] govern contributor terms on external servers. Open platforms including Loculus and Pathoplexus [8], alongside continental sharing frameworks [9], advance pathogen governance, yet regional institutions still benefit from compact, Git-native software unifying versioned metadata, shared binary storage, commit-time biological checks, and revocable institutional custody.

Supporting FAIR interoperability [5] alongside technical mechanisms compatible with CARE-informed governance [10] requires separating software enforcement from institutional ethics processes:

1. **Uniform ingestion validation**: SHA-256 content addressing, offline African Union boundary polygons, and biological screening enforced identically on HTTP commits and raw `git push` hooks.
2. **Institutional custody and controlled sharing**: Operated at https://ubuntubio.org and designed for single-root institutional servers, keeping gated datasets locked until requesters authenticate via ORCID, accept a repository DUA, and obtain custodian approval following external ethics review.
3. **Accountability enablers**: Optional attribution and `benefit_sharing` metadata fields, suppression of the Git LFS batch-negotiation object-existence oracle for inaccessible private blobs, and tamper-evident audit exports with RS256 checkpoints verifiable against off-host custodian anchors. Collective authority and benefit sharing remain the responsibility of institutional ethics committees outside software.

This paper describes the design, security boundaries, crash-recovery guarantees, and single-node evaluation of **UbuntuBio** (v1.0.0), with the hosted portal accessible at https://ubuntubio.org and the client CLI distributed via PyPI (ubuntu-bio).

## 2 Background and Related Systems

Table 1 summarizes six repository architectures across built-in governance custody, binary deduplication, pre-commit host checks, local search, and deployment footprint, distinguishing native capabilities from external pipelines.

**Table 1:** Native capabilities of genomic sequence repositories and surveillance platforms.

| System | Governance Model | Binary Deduplication | Pre-Commit Host Check | Local Metadata Search | Hosting Footprint |
|---|---|---|---|---|---|
| ENA / INSDC [4] | Central public archive | Archive-internal | Post-upload checks | Global portal | National cluster |
| EGA [6] | DAC committee (human) | Archive-internal | Human-cohort scope | Central catalog | Consortium cluster |
| GISAID [7] | Central DUA portal | Platform-internal | Curation review | Portal queries | External central |
| Loculus / Patho-plexus [8] | Open or embargoed | Object store / S3 | External workflows | Backend SQL / API | Multi-service K8s / Docker |
| Git + standard Git LFS | Repo ACLs only | Repo-scoped OID | None built in | None built in | Single host + LFS |
| **UbuntuBio (v1.0.0)** | **Institutional Public / Gated / Private** | **Visibility-scoped SHA-256 CAS** | **Taxon 9606 + NE55 + 9,350-probe** | **Embedded SQLite WAL + FTS5 BM25** | **Single-root directory** |

### Host Provenance vs. Host Sequences

Surveillance datasets record MIxS-derived clinical fields [11] paired with NCBI Taxonomy IDs [12]. Clinical microbial swabs routinely specify `host: Homo sapiens` to document sample provenance while sequence reads represent the pathogen. Tools such as Hostile [13] deplete host reads upstream, yet repositories rarely screen uploaded streams at commit time. UbuntuBio separates provenance from payload: it permits samples declaring `host: Homo sapiens`, rejects human organism records (Taxon ID `9606`), and runs a bounded commit-time screen for targeted human-associated markers. We explicitly treat this screen as a targeted safety check, not a substitute for upstream full-file host depletion.

### Content-Addressed Storage and Open Pathogen Platforms

AllTheBacteria [14], Genome Warehouse [15], and OpenGWAS [16] improve reuse through uniform packaging, while Hugging Face’s Xet backend popularized GearHash sub-file chunking [17]. UbuntuBio `v1.0.0` uses whole-file SHA-256 deduplication within standard Git LFS workflows to minimize operational dependencies, leaving GearHash chunking to future work.

## 3 System Architecture

UbuntuBio keeps mutable curation metadata (cards, TSVs, and approximately 130-byte LFS pointers) inside bare Git repositories while placing sequence binaries exclusively in CAS, routing HTTP API commits, CLI uploads, and raw Git pushes through a single policy gate (Figure 1).

**Figure 1:**
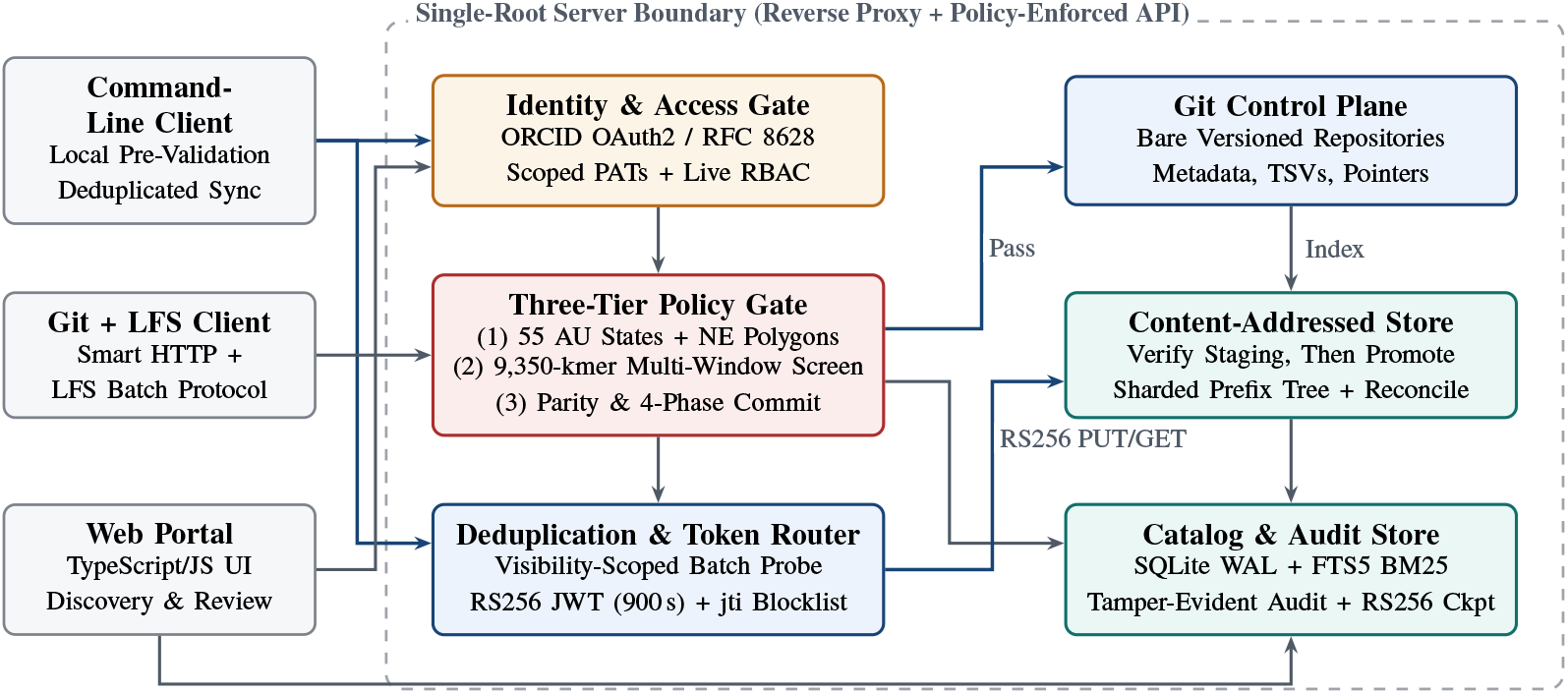
UbuntuBio system architecture. Researchers interact via the web portal (`ubuntubio.org`), MIT-licensed PyPI CLI (`ubuntu-bio`), or standard Git/LFS clients. Submissions traverse identity checks, the shared three-tier policy gate (identical on HTTP commits and Git `pre-receive` hooks), and the four-phase recoverable commit pipeline.

### 3.1 Single-Root Deployment and Four-Phase Recoverable Commit Pipeline

UbuntuBio deploys behind a TLS reverse proxy (Caddy in `deploy/compose.yaml`) as a FastAPI service backed by one directory root holding four stores: (1) bare Git repositories containing dataset cards, sample TSVs, remote file tables, and lightweight Git LFS pointers; (2) a two-level prefix-sharded CAS pool (`<aa>/<bb>/<oid>.bin`); (3) an embedded SQLite (`3.50.4`) WAL database with FTS5 indexes; and (4) an ephemeral staging directory (`tmp/`) for streaming SHA-256 verification. Relational access uses SQLAlchemy so heavier multi-writer deployments can migrate to PostgreSQL in `v2`.

Git LFS `PUT` requests stream bytes exclusively into `tmp/<oid>.tmp` before validation. Because filesystem renames, bare Git updates, and SQLite commits cannot span a single distributed transaction, UbuntuBio uses a **four-phase commit pipeline recoverable via cross-plane reconciliation**:

1. **Phase 1 (Validate)**: Run Tier 1–3 policy checks across cards, sample sheets, parity, and sequence streams. Failures abort with HTTP `422` without touching CAS, Git refs, or SQLite.
2. **Phase 2 (Promote CAS)**: Move each staged file from `tmp/` into `xet/` via `os.replace` (`promote_ to_cas_atomic`). Partial batch failures leave remaining files in `tmp/`, Git refs unchanged, and SQLite unchanged while promoted blobs await orphan cleanup.
3. **Phase 3 (Commit Git)**: Write LFS pointers (`oid sha256:…`) and metadata objects into a temporary `GIT_INDEX_FILE`, build tree/commit objects, and atomically update `refs/heads/<branch>` using Git’s compare-and-swap ref lock. Crashes before `update-ref` leave only unreachable loose Git objects and unreferenced CAS blobs; crashes after `update-ref` leave committed pointers protecting every CAS blob while SQLite lacks catalog mappings.
4. **Phase 4 (Commit Catalog and Index)**: Record `XetFileModel` and `LfsObjectModel` rows inside one SQLite transaction, then refresh `datasets_fts` and `samples_fts` via `index_ repository`.

Administrative reconciliation (`reconcile_cas_storage`) and catalog reindexing (`reindex_all_ repositories`) close failures across all four phases: reconciliation scans every bare Git branch head for LFS pointers and size-matched blobs, protects all Git-referenced CAS blobs, backfills missing SQLite links from Git plus CAS, rebuilds stale FTS5 rows, deletes uncommitted CAS orphans past the `orphan_grace_seconds` threshold (HTTP admin default 3,600 s), and runs `git prune` with `--expire=<N>.seconds.ago` across bare repositories to purge unreachable loose Git objects. Repeated reconciliation passes are idempotent.

### Concurrency and Grace-Period Scope

In `v1.0.0`, `reconcile_cas_storage` runs without a repository-level mutex, and concurrent raw `git http-backend` pushes are serialized at reference update rather than across full push lifetimes (while SQLite WAL serializes catalog commits). A single `orphan_grace_seconds` value governs both CAS blob `mtime` checks and `git prune --expire`. Consequently, `git prune` relies on monotonic filesystem timestamps and assumes active commits complete within the grace window; long-running uploads exceeding 3,600 s or non-monotonic clock jumps risk premature orphan cleanup unless pruning is scheduled during maintenance windows. Our fault-injection suite tests exact transition boundaries in-process rather than OS `SIGKILL` or storage power loss.

### 3.2 Visibility-Scoped Suppression of the Git LFS Batch Oracle

Global CAS eliminates duplicate copies when consortia analyze shared isolates, yet naive deduplication leaks object existence: an unprivileged caller could probe whether another organization holds digest ℎ by requesting an upload slot and observing `actions = null`. Under an authenticated Git LFS batch-negotiation threat model, UbuntuBio enforces **batch-response-shape uniformity** via visibility-scoped transfer rules:

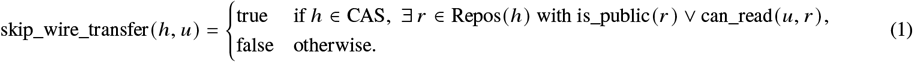

Wire transfer is skipped only if ℎ exists in a public repository or one caller *u* is authorized to read. When ℎ resides solely in inaccessible private or unapproved gated repositories, batch negotiation issues a standard write token and upload URL whose JSON structure matches unknown digests.

During post-upload execution, `PUT …/lfs/oid` streams bytes directly into `tmp/<oid>.tmp`, verifies the SHA-256 digest without querying CAS, and uniformly returns `{“status“: “uploaded”, …}`. In Phase 2, `promote_to_cas_atomic` checks whether `xet/…/<oid>.bin` already exists; if present, it unlinks `tmp/<oid>.tmp`, preserves the single canonical CAS blob on disk, and returns the standard HTTP `201` commit payload without revealing deduplication status. Concurrent `PUT` streams for identical unknown digests share `tmp/<oid>.tmp`, and network timing analysis or storage-quota channels remain outside the claimed security boundary of `v1.0.0`.

### 3.3 Uniform Policy Enforcement Across HTTP Commits and Git Pushes

UbuntuBio `v1.0.0` enforces identical Tier 1–3 rules across REST `/commit` and the server-side Git `pre-receive` hook (`ubuntubio.git.validator_hook`). During raw `git push`, the hook extracts inline sequence blobs directly and resolves LFS pointers against `tmp/<oid>.tmp` or `xet/`; any dangling or un-uploaded LFS sequence pointer is rejected so raw Git pushes cannot bypass validation.

#### Tier 1: Core MIxS-Derived Subset, AU-55 / Outgroup Scope, and Natural Earth Polygons

Repositories provide a YAML front-matter card in README.md (title, public/gated/private tier, domain, license or DUA, optional attribution and benefit_sharing fields, and policy toggles) alongside nine mandatory core MIxS-derived columns in samples.tsv (sample_id, taxon_id, country, collection_date, lat, lon, admin1, host, isolation_source); extended MIxS environmental package columns are stored optionally in extra_fields without claiming full GSC MIxS certification.

Table 2 specifies geographic policy semantics implemented in `validator/models.py`. Under the default `au_strict` policy, both `README.md:countries` and every row in `samples.tsv` must use the 55 AU member state ISO-3166-1 alpha-2 codes (including Western Sahara, `EH`). Under `geo_policy: allow_outgroups` (designed for comparative phylogenetics), `README.md:countries` must still contain at least one AU member state anchor while permitting non-AU ISO-3166-1 alpha-2 country codes (e.g., `US`); non-AU outgroup rows require valid WGS-84 bounds ([−90, 90] × [−180, 180]) without matching an African polygon, whereas AU rows continue full polygon validation.

**Table 2:**
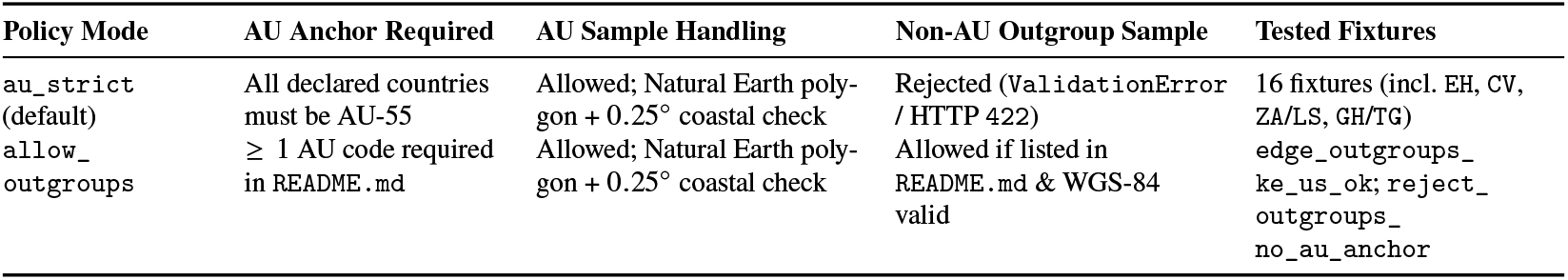
Implemented geographic validation rules by repository geo_policy.

| Policy Mode | AU Anchor Required | AU Sample Handling | Non-AU Outgroup Sample | Tested Fixtures |
|---|---|---|---|---|
| au_strict (default) | All declared countries must be AU-55 | Allowed; Natural Earth poly-gon + 0.25° coastal check | Rejected (ValidationError / HTTP 422) | 16 fixtures (incl. EH, CV, ZA/LS, GH/TG) |
| allow_outgroups | ≥ 1 AU code required in README.md | Allowed; Natural Earth poly-gon + 0.25° coastal check | Allowed if listed in README.md & WGS-84 valid | edge_outgroups_ke_us_ok; reject_outgroups_no_au_anchor |

Coordinates for AU member states are checked in pure Python against quantized multi-polygons from **Natural Earth 1:50m Admin-0 Countries** (`au_boundaries.json.gz`): (1) for each polygon `poly`, `_point_in_polygon` applies even-odd ray casting to verify inclusion inside the exterior ring (`poly`[0]) and exclusion from all interior holes (`poly`[1:]) so enclaves inside holes (e.g., Lesotho within South Africa) cannot pass under the surrounding country’s code; (2) before granting coastal tolerance, all other 54 AU state polygons are checked so interior points in bordering neighbors (e.g., Togo near Ghana’s border) fail immediately; and (3) remaining offshore points pass only when squared Euclidean distance in WGS-84 `(lon, lat)` degrees to an exterior-ring segment (`poly`[0]) satisfies *d* ≤ 0.25^◦^. Because distance is computed in unprojected degrees rather than geodesic meters, 0.25^◦^ is an angular approximation spanning 22.0–27.8 km east–west and 27.8 km north–south across Africa ([−35.5^◦^, 37.5^◦^]). These boundaries act solely as technical curation filters rather than legal adjudications.

#### Tier 2: Taxon 9606 Ban, QC Checks, and Server-Keyed Targeted Human-Marker Screen

Samples declaring NCBI Taxon ID `9606` are rejected. On FASTQ, FASTA, and BAM streams (raw, gzip, or BGZF), UbuntuBio validates syntax and sequence/quality length parity, warns on legacy `Phred+64` quality or > 15.0% `N` bases, and rejects headers matching human assembly tokens (*Homo sapiens*, GRCh37/38, hg19/38, `NC_0000*`). **CRAM streams undergo magic-byte recognition only** (`CRAM`): read syntax, contigs, and human 25-mers are not validated on CRAM inputs.

For FASTQ/FASTA/BAM read screening, UbuntuBio matches 25-mers against a **targeted panel of 9,350 human-associated probes** (`human_kmers.txt.gz`). The panel was built by tiling human mitochondrial DNA (`NC_012920.1`), 18S rRNA (`NR_003286.4`), 28S rRNA (`NR_003287.4`), and high-copy AluY repeats (`U14574.1`)—yielding 5,861 positive target tiles—and subtracting exact 25-mers shared with selected bacterial, viral, mitochondrial, *Plasmodium*, and *Trypanosoma* genomes. Fungal rRNA references were omitted during `v1.0.0` subtraction, accounting for the conserved 18S rDNA cross-match observed on *Candida albicans*. We distinguish: *human-associated marker detection* (25-mer matches against this repeat/rDNA panel), *human-specific sequence detection* (unique non-conserved human loci), and *verified human DNA contamination* (full reference alignment/depletion).

In `multi_window` mode (`max_reads = 2,000`, `max_scan_records = 250,000`), scanning caps total records inspected at 250,000: records 1–1, 000 are checked sequentially, and Algorithm R reservoir sampling [18] selects 1,000 reads uniformly without replacement from records 1, 001–*N*_scan_ where *N*_scan_ = min(*N*, 250, 000). Consequently: (1) files with *N* ≤ 2, 000 reads have every read inspected deterministically without reservoir eviction; and (2) in files with *N* > 250, 000 reads, contamination confined past record 250,000 lies outside the bounded sampling frame. The PRNG is seeded via HMAC-SHA256 from a server secret and stream path; re-uploading identical LFS blobs hashes to the same staged path within a server lifecycle, so submission rate-limiting relies on the TLS reverse proxy. Stewards configure `human_screen_policy` as `warn` (default) or `strict_reject`.

#### Tier 3: Metadata-to-Tree Parity

Sequencing libraries declared in README.md must match committed files or remote_files.tsv rows with zero missing or orphaned sequence paths.

### 3.4 Gated Custody, Device Authentication, Tokens, and Audit Anchors

Researchers authenticate via ORCID OAuth 2.0 Authorization Code (/api/auth/orcid/authorize and /callback) or UbuntuBio’s server-hosted RFC 8628 Device Authorization endpoints [19] (/api/auth/device/code, /verify, /token): because ORCID itself is not used as a direct RFC 8628 endpoint, the CLI requests a 10-minute UBIO-XXXX-XX code from UbuntuBio, the researcher signs into the web portal with ORCID, and approving /device mints a single-use 90-day CLI PAT bound to their ORCID UserModel.

Authority over gated datasets rests with each repository’s designated custodians and organization administrators (Figure 2). Requesters record an **authenticated DUA acceptance event** in `dua_acceptances` (binding their ORCID-authenticated `user_id`, `repo_id`, `dua_version`, and UTC timestamp inside the hash-chained audit log) rather than a client-side asymmetric cryptographic signature; requests without DUA acceptance return HTTP `400`, and pending requests return HTTP `403`. Once approved, 900-second capability JWTs [20, 21] are issued with claims `iss` (server API URL), `aud = “openxet“`, `sub` (ORCID iD), `iat`, `nbf`, `exp`, unique `jti`, `repo`, `action`, and `oid`. Every CAS download verifies: (1) RS256 signature under a strict `algorithms=[“RS256”]` allow-list against active or retired `*.pub.pem` keys exposed at `/.well-known/jwks.json` [22] across 2048-bit rotation; (2) absence from the `RevokedCasTokenModel jti` blocklist table; and (3) live `AuthorizationService` permission lookup so custodian revocation rejects unexpired JWTs immediately (HTTP `403`).

**Figure 2:**
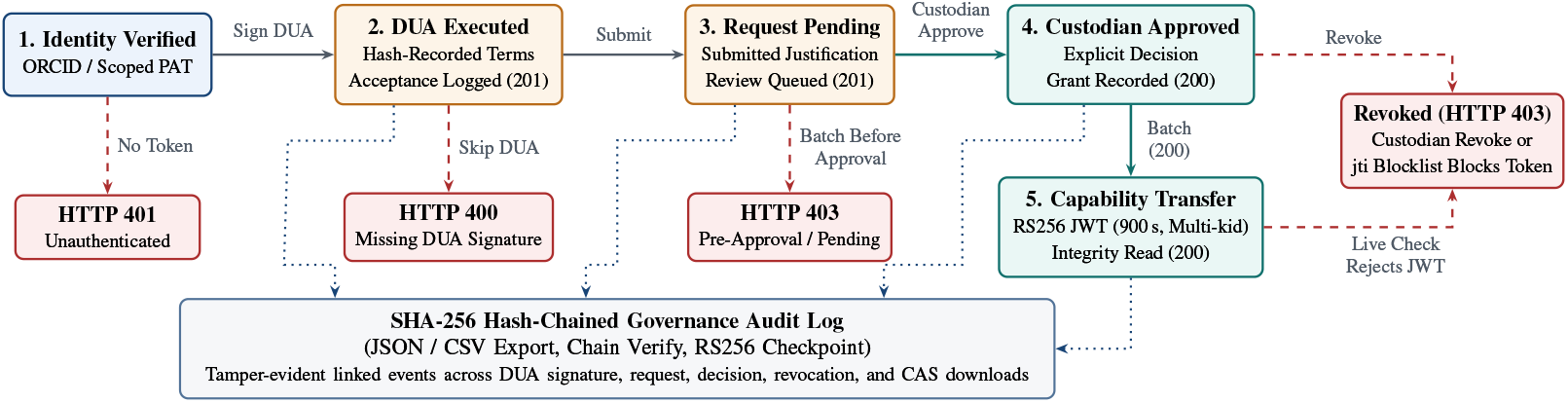
Gated institutional custody state machine. Unauthenticated batch requests return HTTP 401; requests missing DUA acceptance return HTTP 400; pending requests receive HTTP 403. Approval by designated custodians unlocks 900-second RS256 capability tokens checked against JWKS keys, a `jti` blocklist, and live database permissions. RS256 audit checkpoints establish tamper evidence relative to independently retained custodian anchors.

Governance and gated download events write sequentially to a **SHA-256 hash-chained `audit_log`**, and custodians can fetch an RS256-signed audit checkpoint JWT binding `head_hash` and `entries_ checked`. *Operational trust model*: an active server private key cannot stop a rogue administrator controlling both S QLite and disk keys from rewriting rows and re-signing a fresh checkpoint. Therefore, institutional compliance officers or consortium oversight partners archive periodic checkpoints off-host (e.g., on release/daily schedules or on append-only WORM storage) via `ubio audit checkpoint --save <ledger.jsonl>` and verify exports against those external anchors via `--verify-against`.

### 3.5 Visibility-Scoped SQLite WAL + FTS5 Catalog

Metadata indexes into SQLite (`3.50.4`) WAL FTS5 virtual tables (`datasets_fts` and `samples_fts`). Query strings are sanitized into prefix-wildcard tokens joined by FTS5 implicit `AND`. Dataset search applies weighted BM25 [23] across `slug`, `title`, `description`, `countries`, `genus`, `family`, and `benefit_sharing` with weights (10.0, 5.0, 1.0, 3.0, 3.0, 2.0, 1.0), while sample search uses unweighted BM25 and filters private and unapproved entries against caller permissions.

## 4 Implementation and Client Ecosystem

UbuntuBio `v1.0.0` comprises: (1) the Python 3.12+ FastAPI server backend verified across 96 unit and integration tests (75% total line coverage, including real `validator_hook` Git subprocess tests, token revocation, audit rewrite verification, and CAS reconciliation); (2) the MIT-licensed `ubuntu-bio` Python CLI on PyPI (`pip install ubuntu-bio`), verified across 67 tests (73% line coverage); and (3) the TypeScript/Next.js web portal at https://ubuntubio.org. A packaged bundle of server modules, deployment manifests, and reproducible evaluation benchmarks is maintained privately for institutional deployment and made available upon reasonable editorial request.

## 5 Evaluation

All benchmarks ran on an **Apple M4** workstation (10 logical cores, 16 GiB RAM, Darwin 27.2.0, local APFS NVMe SSD) using Python 3.12.13 and SQLite 3.50.4 inside isolated temporary workspaces (benchmarks/benchmark_paper.py), representing single-node execution.

### 5.1 Policy Gate Parity Across 18 Boundary Fixtures

To verify consistent enforcement across ingestion paths, we tested **18 policy fixtures** (Appendix A): three baseline curated cohorts, six positive boundary cases (`EH`, `CV`, nearshore Ghana ≤ 0.25^◦^, `allow_outgroups` anchored by Kenya, `Phred+64` warning, and `human_screen_policy: warn`), three baseline negative fixtures (Taxon `9606`, non-AU `US`, out-of-polygon Kenya coordinate), and six negative boundary fixtures (Lesotho enclave mislabeled `ZA`, Togo border point 0.02^◦^ east of Ghana rejected by neighbor check, `allow_outgroups` lacking an AU anchor, `strict_reject` k-mer block, FASTQ seq/qual mismatch, and Tier 3 missing declared file). Python validation, Git `pre-receive`, and HTTP `/commit` agreed on **18/18** cases (9 positive HTTP `201` commits in 343.12–901.77 ms; 9 negative HTTP `422` rejections in 207.01–259.29 ms). These deterministic fixtures verify explicit rule boundaries rather than random population generalization.

### 5.2 Targeted Human-Marker Screening: Empirical Cross-Match and Statistical Sensitivity

We evaluated the 9,350-probe panel across four separated experiments (Tables 3 and 4):

1. **Panel construction check vs. 22 RefSeq target records (19 organisms)**: Checking the panel back against its 5,861 positive target tiles yields 100.0% in-sample coverage by construction. On an independent simulated test across **280,157 150 bp reads** from **22 NCBI RefSeq sequence records spanning 19 distinct organisms/loci**—7 bacterial genomes (7 seqs), **7 parasite sequences across 4 named organisms** (*P. falciparum* chr1, apicoplast, mitochondrion, 18S rRNA; *T. brucei* chr1; *L. major* chr1; *S. mansoni* mitochondrion), 1 fungal chromosome (*Candida albicans* chrR, `NC_032096`), and 7 priority viruses (7 seqs)—the empirical cross-match rate was **0.000 across 21/22 sequences**. *C. albicans* chrR matched 17/15,000 reads (0.113%) at conserved eukaryotic 18S rDNA positions (aggregate cross-match rate: 6.1×10^−5^, 95% Wilson CI [3.8×10^−5^, 9.7×10^−5^]). This bounded set measures empirical cross-matching on selected reference taxa rather than clinical specificity across all pathogen diversity.
2. **Independent human sequencing sensitivity**: Excluded from panel design, an independent *Homo sapiens* WGS run (`SRR30459529`, 6.39 MiB) yielded **1,734/180,114 marker hits (0.96% density)** in full scan and **12/2,000 hits** in one `multi_window` pass. Pure single-copy human *ACTB* mRNA (`NM_001101.5`) triggered human header rejection while returning 0 k-mer hits, showing that repeat/rDNA probes detect genomic carryover rather than isolated coding exons.
3. **Real African ENA primate carryover**: Across 174,571 sampled reads from five African *E. coli* ENA runs, four files had 0 hits and `SRR11015023_1.fastq.gz` flagged 1 sampled hit (5/114,852 full-scan reads). Checking ENA `PRJNA604701` traced this run to *Chlorocebus sabaeus* (African green monkey) stool in The Gambia; flagged reads shared 5–37 exact 25-mers with primate Alu-Sx (`U14574.1`) and 0 with *E. coli* K-12, demonstrating *human-associated primate repeat detection* rather than verified *Homo sapiens* contamination.
4. **Separate uniform vs. interior statistical expectations and 0.01% seed audit**: Across 30 seeds per contamination fraction (*p* ∈ [10^−4^, 5 × 10^−2^], 50,000 *E. coli* reads; Figure 3(a) and Table 4), uniform spike-ins span reads 0–49, 999 (*P*_uni_(detect) ≈ 1 − (1 − *p*)^2000^), whereas interior spike-ins (*k*_spike_ = max(1, round(50, 000 *p*))) are placed uniformly without replacement in reads 10, 000–49, 999 (`lo_idx = 10,000`). Because reads 0–999 are always clean, interior detection relies solely on the 1,000 reservoir-sampled reads drawn across the 49,000 post-prefix reads, following *P*_int_(detect) ≈ 1 − (1 − *p*)^1000^ (exact hypergeometric-like conditional 1 − (1 − *k*_spike_/49, 000)^1000^ = 40.0%, 64.0%, 99.4% closely matches observed 12/30 = 40.0%, 19/30 = 63.3%, 30/30 = 100.0%). At *p* ≥ 0.5%, `multi_window` detected 30/30 seeds in both regimes (95% Wilson CI [24]: [0.886, 1.000]) in 56.2–56.9 ms, while prefix `head` scanning missed 0/30 interior spikes.

**Figure 3:**
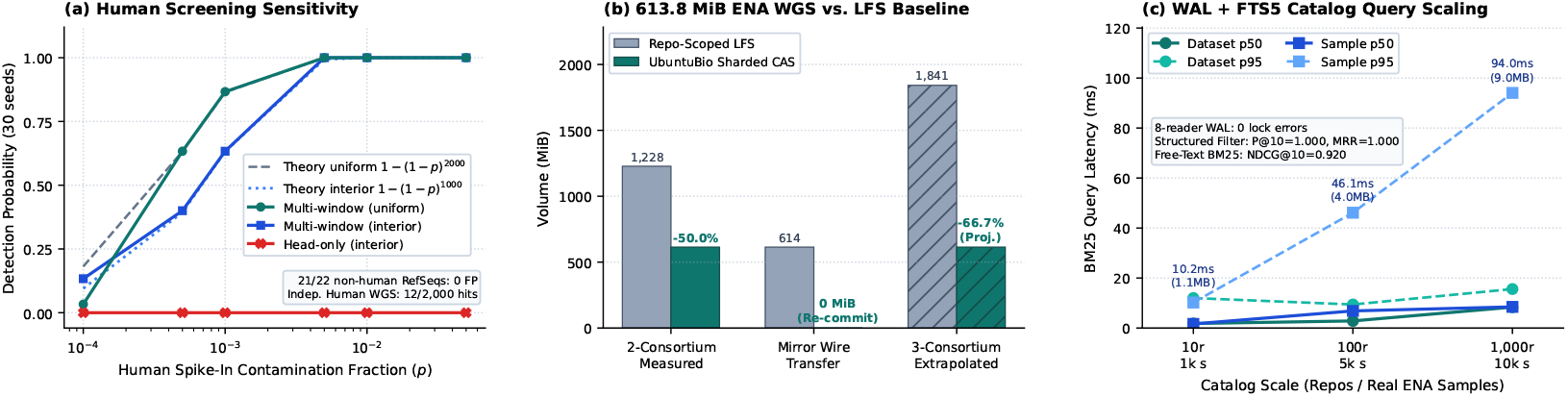
Empirical evaluation on single-node Apple M4 hardware. (a) 30-seed spike-in curves against separate uniform 1 − (1 − *p*)^2000^ and interior 1 − (1 − *p*)^1000^ binomial expectations, with 0/265,157 false cross-matches across 21/22 non-human RefSeq sequences. (b) Component-accounted 2-consortium storage (measured) vs. 3-consortium analytical extrapolation (hatched). (c) SQLite WAL + FTS5 scaling across 10,000 real ENA biosamples.

**Table 3:** Empirical cross-match rate of the 9,350-probe panel across 22 RefSeq sequences (19 organisms/loci, 280,157 reads), independent human WGS, and 5 African ENA *E. coli* runs.

| **Dataset / RefSeq Target** | **Sequences (Organisms)** | **Reads** | **Hits** | **Rate** | **95% Wilson CI** |
|---|---|---|---|---|---|
| Bacterial Pathogens (*E. coli*, *K. pneu.*, *M. tb*, *V. chol.*, *S. aur.*, *S. ent.*, *A. baum.*) | 7 seqs (7 orgs) | 105,000 | 0 | 0.000 | [0.0, 3.7 × 10<sup>−5</sup>] |
| Parasite Loci (*P. falc.* chr1/api/mt/18S, *T. brucei*, *L. major*, *S. mansoni* mt) | 7 seqs (4 orgs) | 72,027 | 0 | 0.000 | [0.0, 5.3 × 10<sup>−5</sup>] |
| Priority Viruses (Mpox, SARS-CoV-2, Lassa, Ebola, RVFV, Yellow fever, HIV-1) | 7 seqs (7 orgs) | 88,130 | 0 | 0.000 | [0.0, 4.4 × 10<sup>−5</sup>] |
| *Candida albicans* SC5314 chrR (NC_032096; conserved 18S rDNA) | 1 seq (1 org) | 15,000 | 17 | 0.001133 | [7.08 × 10<sup>−4</sup>, 1.81 × 10<sup>−3</sup>] |
| Real African ENA *E. coli* (5 runs; 1 *Chlorocebus sabaeus* Alu hit) | Clinical WGS | 174,571 | 1 | 6.0 × 10<sup>−6</sup> | [1.0 × 10<sup>−6</sup>, 3.2 × 10<sup>−5</sup>] |
| Independent Human WGS (SRR30459529; full / 2k multi_window) | *Homo sapiens* | 180,114 | 1,734 | 0.0096 | (12/2,000 in 2k draw) |

**Table 4:** 30-seed spike-in detection counts (95% Wilson CIs) against regime-specific binomial expectations: *P*_uni_ = 1 − (1 − *p*)^2000^ for uniform placement and *P*_int_ = 1 − (1 − *p*)^1000^ for interior reservoir placement (≥ read 10,000).

| **Fraction *p*** | **Unif. Binom. (2k)** | **Uniform MW (CI)** | **Int. Binom. (1k)** | **Interior MW (CI)** | **Interior Head** | **MW Latency** |
|---|---|---|---|---|---|---|
| 0.0001 (0.01%) | 18.13% | 1/30 [0.006, 0.167] | 9.52% | 4/30 [0.053, 0.297] | 0/30 | 56.56 ms |
| 0.0005 (0.05%) | 63.22% | 19/30 [0.455, 0.781] | 39.35% | 12/30 [0.246, 0.577] | 0/30 | 56.89 ms |
| 0.0010 (0.10%) | 86.48% | 26/30 [0.703, 0.947] | 63.23% | 19/30 [0.455, 0.781] | 0/30 | 56.71 ms |
| 0.0050 (0.50%) | 99.996% | 30/30 [0.886, 1.000] | 99.33% | 30/30 [0.886, 1.000] | 0/30 | 56.56 ms |
| 0.0100 (1.00%) | 100.0% | 30/30 [0.886, 1.000] | 100.0% | 30/30 [0.886, 1.000] | 0/30 | 56.20 ms |
| 0.0500 (5.00%) | 100.0% | 30/30 [0.886, 1.000] | 100.0% | 30/30 [0.886, 1.000] | 0/30 | 56.82 ms |

Investigating the low uniform *p* = 0.0001 count (1/30 vs. 18.13% expected, ≈ 5.4/30) at seed granularity (Appendix D) showed zero bugs: across the 30 seeded trials (5 spikes each), 0/30 happened to place a spike inside reads 0–999 (*P* = 0.9039^30^ = 4.8%), 4/30 placed spikes in reads 1, 000–1, 999 (all caught deterministically by `head`), and trial 4 (`seed=10103`, read `5277`) landed in the 1,000-read reservoir (one-sided Binomial *p* = 0.0185). A decoupled **200-seed independent verification** at *p* = 0.0001 confirmed unbiased convergence: `multi_window` detected **38/200 (19.0%, 95% CI** [0.142, 0.250]) vs. theoretical 18.13% (and `head` detected 44/200, 22.0%).

### 5.3 613.83 MiB Storage Component Accounting, Local Transfer, and Batch Uniformity

We evaluated a **613.83 MiB (643,646,879 B) Nigerian *E. coli* paired-end WGS cohort**, comprising runs `SRR9681899_1/2`, `SRR9822635_1/2`, and `SRR9822643_1/2` plus manifests (8 files), against a repo-scoped baseline (two independent repository binary directories containing identical files on the same APFS NVMe filesystem; Table 5 and Figure 3(b)):

1. **Component accounting vs. analytical projection**: Across two measured repositories, UbuntuBio stored **613.83 MiB (643,649,025 B)** in `xet/` CAS, **0.05 MiB (56,964 B)** in bare Git control planes, and **0 B** remaining in `tmp/` (613.88 MiB total vs. 1,227.66 MiB repo-scoped payload copies; **50.0% sequence storage reduction**). Primary commit took 3,081.47 ms (199.20 MiB/s); repeat mirror commit avoided wire transfer for all 8 accessible objects (**0 B transferred**, 2,539.93 ms CPU validation). Figure 3(b) and Table 5 explicitly separate the measured two-repository result from a three-consortium analytical extrapolation (3 × 613.83 = 1, 841.49 MiB vs. 613.83 MiB, −66.7%) assuming whole-file sharing across a third consortium.
2. **Local in-process vs. WAN transfer**: Fetching 492.5 MiB across 4 concurrent local FastAPI ASGI readers completed in **410.33 ms (1,200.27 MiB/s)**; raw NVMe worker reads under `F_NOCACHE` (18.30 ms) vs. warm page cache (17.47 ms) capture local controller/memory bandwidth rather than African institutional WAN throughput (where 492.5 MiB across 10– 100 Mbps links requires 41–413 s).
3. **Batch response-shape uniformity**: Combining the 93.66 MiB ENA cohort with two private commits (`priv-alpha` and `priv-beta`, 1,590,821 B each) yielded 199,607,808 B logical data vs. 99,803,904 B physical CAS (1 physical blob). Running 60 interleaved batch probes from `priv-beta` checking `priv-alpha`’s private digest against an unknown OID produced **100% identical HTTP 200 JSON shapes**, with private-hit mean 24.292 ms (median 24.103 ms, IQR 0.417 ms, p95 24.558 ms) vs. unknown-OID mean 24.276 ms (median 23.424 ms, IQR 0.447 ms, p95 25.486 ms; paired delta 0.017 ms, 95% CI [−1.084, 1.117] ms). While this local CI includes zero, it proves batch-response-shape uniformity rather than timing-channel indistinguishability.

**Table 5:** Single-root storage component accounting on 613.83 MiB Nigerian *E. coli* WGS runs.

| **Scenario** | **Basis** | **Repo-Scoped LFS** | **CAS** (xet/) | **Git Control** | tmp/ | **Sequence Savings** |
|---|---|---|---|---|---|---|
| 2-Consortium Repositories | Measured | 1,227.66 MiB | 613.83 MiB | 0.05 MiB | 0 B | 50.0% |
| Mirror Re-commit Wire | Measured | 613.83 MiB | 0 B wire | — | 0 B | 100.0% wire |
| 3-Consortium Sharing | Analytical Ext. | 1,841.49 MiB | 613.83 MiB | — | — | 66.7% (proj.) |

### 5.4 SQLite WAL Scaling, Structured-Filter Correctness, and Free-Text BM25 Ranking

Seeding SQLite WAL catalogs from **10,000 real African *E. coli* biosamples** across three tiers—10 repos / 1,000 samples, 100 repos / 5,000 samples, and 1,000 repos / 10,000 samples (80% `public`, 10% `gated`, 10% `private`)—produced databases of 1.05 MiB, 4.05 MiB, and 8.98 MiB with online `sqlite3` backups finishing in 1.84 ms, 6.43 ms, and 12.08 ms. Over 100 single-reader iterations per index (measured without a separate untimed warm-up pass), median latencies were 1.799, 2.834, and 8.346 ms for dataset search (p95: 11.989, 9.409, 15.590 ms) and 1.637, 6.842, and 8.454 ms for sample search (p95: 10.190, 46.118, 94.040 ms; Figure 3(c)). Under 8 concurrent async reader tasks (15 queries/task, 120 total queries) running alongside 1 active writer (5 commits), **aggregate reader throughput** reached 220.44, 65.57, and 39.15 QPS with **0 SQLite lock errors**.

#### 5.4.1 Structured-Filter and Access-Control Correctness (27 Queries, 10,000 Samples)

At 1,000 repos / 10,000 samples, we evaluated 24 positive structured-filter/token queries and 3 negative zero-hit checks against SQL ground-truth rules over visible records (10% private excluded; Appendix B). Because these queries test matching predicates (e.g., `country=NG`, `source LIKE %stool%`), perfect top-K accuracy verifies filter fidelity, `limit=20` pagination, and private-record isolation rather than semantic BM25 ranking. With Recall@10 = |relevant in top 10|/|all relevant| upper-bounded at 10/|Rel.| when |Rel.| > 10, UbuntuBio achieved macro P@5 = 1.000, macro P@10 = 1.000, macro Recall@10 = 0.3157, 3/3 negative zero-hit checks, and **0 private leaks**.

#### 5.4.2 Independent Free-Text Weighted BM25 Ranking (Experiment 5b)

To evaluate BM25 ranking separately from SQL filters, we built a 23-repository multi-pathogen benchmark (`bacteria`, `parasite`, `virus`, `metagenome`; 17 `public`, 5 `gated`, 1 `private`) and issued **6 filter-free multi-term queries** evaluated against pre-specified graded domain relevance labels (rel = 2 primary target cohort, rel = 1 secondary regional mention, rel = 0 distractor matching query tokens in `benefit_sharing` or comparator notes; Appendix D, Table 10). Weighted FTS5 BM25 achieved **macro P@5** = 0.6945, **macro Recall@10** = 0.9167, **macro MRR (primary rel** = 2) = 1.000, **macro graded NDCG@10** = 0.9200, and **0 private leaks**. Primary cohorts consistently outranked `benefit_sharing` distractors (NDCG@10 = 1.000 on *Malawi bloodstream*, *Ghana malaria resistance*, and *Kenya cholera outbreak*), while revealing realistic bag-of-words boundaries: an Ethiopian description distractor interleaved between two South African TB cohorts (NDCG@10 = 0.9197) and `senegal pediatric diarrhea` missed a synonym cohort titled *Childhood Gastroenteritis* (Recall@10 = 0.500, NDCG@10 = 0.6131).

### 5.5 Gated Custody, Anchor Rewrite Detection, and 7-Transition Crash Matrix

Gated probes verified HTTP `401`, `400`, `403` (pending, mid-session custodian revocation on an unexpired 900 s JWT, and admin `jti` blocklist), and `200` SHA-256 download, while key rotation preserved historical `ubuntubio-cas-key-1` alongside `ubuntubio-cas-key-1794833404-dfc5a8` in JWKS. Editing SQL row `5a0f36d5-…` broke `/audit-log/verify` and blocked checkpoint signing (`409`); when a simulated rogue administrator recomputed the entire SHA-256 chain from row 1 and signed a fresh `200` checkpoint, comparison against the retained off-host anchor detected the rewrite. Across all 7 injected phase-transition faults (Table 6, running in-process teardown without OS `SIGKILL`), `reconcile_cas_storage` (including `git prune`) recovered 7/7 transitions cleanly, confirmed second-pass idempotency, and preserved fresh blobs inside the 3,600 s grace window.

**Table 6:** 7-transition crash-injection matrix (T1–T7), idempotency, grace preservation, and security checks.

| Scenario / Transition | Injected Fault / Security Probe | Post-Reconcile / Observed Outcome |
|---|---|---|
| Security & Key Rotation | Mid-session revoke; jti block; JWKS rotation | Live HTTP 403 immediately; 2 active/retired JWKS kids |
| Audit & Rogue Rewrite | Tamper SQL row; recompute chain + re-sign | /verify catches ID; ckpt 200 to 409; anchor catches rewrite |
| T1: Phase 1 Abort | Taxon 9606 commit rejected before Phase 2 | HTTP 422; 0 CAS promoted; Git HEAD & DB un-changed |
| T2: Phase 2 Partial | 2-file rename crash (File A moved, File B failed) | Orphan File A pruned; Git ref & prior blobs intact |
| T3: Phase 2 Complete | Complete CAS promotion, crash before Git | Aged unreferenced CAS orphan pruned via HTTP reconcile |
| T4: Phase 3 Mid-Git | CAS + loose Git blob written before update-ref | Branch intact; reconcile git prune deletes loose blob |
| T5: Phase 3 Post-Git | update-ref done, DB teardown before SQLite + FTS5 | Reconcile shields 4 blobs, repairs 8 DB links, rebuilds FTS5 |
| T6: Phase 4 Post-DB | SQLite committed, crash before FTS5 index | reindex rebuilds datasets_fts cleanly from Git+DB |
| T7: Mixed + Idempotency | Orphan + dropped DB/FTS5 + repeat pass + active blob | Restores 8 links + FTS5; 2nd pass no-op; 3,600 s blob kept |

## 6 Limitations and Roadmap

Single-node evaluation highlights six boundaries in v1.0.0:

1. **Whole-file vs. chunk deduplication**: SHA-256 deduplicates identical files; GearHash sub-file chunking [17] is future work.
2. **Targeted markers vs. full decontamination**: The 9,350-probe panel hits 0.96% of human WGS reads while cross-matching *C. albicans* 18S rDNA (0.113%) and primate Alu motifs and missing single-copy exons; upstream depletion [13] remains required, with fungal rDNA subtraction planned for `v1.1`. Reads beyond record 250,000 sit outside `multi_window`.
3. **CAS timing and quota channels**: Batch response-shape uniformity does not prevent network timing analysis or quota channels, and concurrent identical `PUT` streams share staging paths.
4. **Reconciliation locks and anchor retention**: `reconcile_cas_storage` uses a single `orphan_ grace_seconds` window for CAS and `git prune` without a repo mutex, assuming off-host anchor retention by stewards.
5. **CRAM parsing and single-node concurrency**: CRAM receives magic-byte recognition only, while SQLite WAL targets campus workloads (39–220 aggregate reader QPS across 8 tasks), with PostgreSQL migration supported in `v2`.
6. **Non-human scope and ethical governance**: Banning Taxon `9606` excludes human host genetics cohorts (better served by EGA [6]), while community benefit governance remains with institutional ethics boards outside software.

## 7 Conclusion

UbuntuBio is a Git-native, policy-enforced genomic repository designed for institutional data custody, controlled sharing, and reproducible workflows across regional pathogen research centers. Combining prefix-sharded CAS with Git LFS pointers, 55-state Natural Earth boundaries plus anchored outgroup support, uniform REST and Git `pre-receive` screening, revocable RS256 tokens, externally anchorable audit checkpoints, and four-phase reconciliation, `v1.0.0` halves two-repository binary storage on 613.83 MiB real WGS cohorts while keeping single-root institutional deployment compact.

## Data and Software Availability

The hosted UbuntuBio portal is accessible at https://ubuntubio.org and the MIT-licensed `ubuntu-bio` CLI is publicly distributed on PyPI (`pip install ubuntu-bio`). Server modules (`server/`), web portal components (`web/`), single-root deployment manifests (`deploy/compose.yaml`), and evaluation harnesses (`benchmarks/`) are maintained under institutional stewardship at WACCBIP, University of Ghana, and are available from the corresponding author upon reasonable request; a permanent archival software snapshot and DOI will be deposited upon peer-reviewed publication. Bundled static validation artifacts carry SHA-256 digests `864ffd73a4182e6b7c7d66ac95f46380 f1b630db701fd5a01b0a0e1173ca6232` (`au_boundaries.json.gz`) and `6fcbd7fdcda9920c899a12 883d1d3ed46e8ceb2ab3209e5f1c9d5f8b6890964f` (`human_kmers.txt.gz`). Complete benchmark outputs and Figure 3 are reproduced from the benchmark package via `server/.venv/bin/python benchmarks/benchmark_paper.py --json-out paper/figures/bench_results.json` and `uv run --with matplotlib --with numpy python paper/figures/make_fig3.py`. All reference sequence accessions and evaluation artifacts are listed in Appendices A–D.

## Acknowledgements

This work was developed by the pioneer class of bioinformatics students (Class of 2027) at the West African Centre for Cell Biology of Infectious Pathogens (WACCBIP), University of Ghana. We thank the faculty, mentors, and bioinformatics and pathogen genomics teams at WACCBIP for their guidance and feedback on sample curation, metadata standards, and institutional access policies.

## Author Contributions

All twenty authors contributed equally to this work, encompassing software architecture, policy-gate implementation, benchmarking, domain validation specification, metadata curation standards, governance policy design, and manuscript preparation, review, and approval.

## Ethics and Data-Use Statement

No human participants or identifiable clinical records were enrolled; all evaluated sequencing benchmarks use publicly archived ENA and NCBI RefSeq records and UbuntuBio enforces strict exclusion of human organism records (Taxon ID 9606).

## A Policy-Gate Boundary Fixtures (18 Cases)

Table 7 records all 18 boundary policy fixtures evaluated identically across Python validation, Git pre-receive, and HTTP /commit.

**Table 7:** Complete 18 policy-gate fixtures, cross-plane agreement, HTTP status codes, and end-to-end latencies.

| **Fixture ID** | **Boundary / Rule Tested** | **Expected** | **Hook / HTTP** | **Code** | **Lat. (ms)** |
|---|---|---|---|---|---|
| malawi-amr-ecoli | Curated public cohort (6 files, MW) | Pass | Pass / Pass | 201 | 901.77 |
| rwanda-mpox-surveillance | Curated public viral cohort (8 files, RW) | Pass | Pass / Pass | 201 | 676.18 |
| kenya-cholera-gated | Curated gated cohort + DUA (7 files, KE) | Pass | Pass / Pass | 201 | 615.39 |
| edge_western_sahara_eh | AU-55 inclusion: Western Sahara (EH) | Pass | Pass / Pass | 201 | 373.20 |
| edge_island_cape_verde | Island state geometry: Cabo Verde (CV) | Pass | Pass / Pass | 201 | 359.94 |
| edge_coastal_buffer_gh | Offshore coastal point ≤ 0.25° from GH | Pass | Pass / Pass | 201 | 359.31 |
| edge_outgroups_ke_us_ok | allow_outgroups (KE anchor + US WGS-84) | Pass | Pass / Pass | 201 | 364.00 |
| edge_phred64_warning_ok | Legacy Phred+64 encoding warning allowed | Pass | Pass / Pass | 201 | 374.42 |
| edge_human_warn_policy_ok | human_screen_policy: warn audit pass | Pass | Pass / Pass | 201 | 343.12 |
| reject_human_9606 | Taxon 9606 + GRCh38 header rejection | Reject | Reject / Reject | 422 | 257.61 |
| reject_non_au_country | Non-AU country (US) under au_strict | Reject | Reject / Reject | 422 | 217.23 |
| reject_invalid_lat_lon | Declared KE coord 58.05° outside polygon | Reject | Reject / Reject | 422 | 207.01 |
| reject_enclave_lesotho_as_za | Lesotho enclave point mislabeled as ZA | Reject | Reject / Reject | 422 | 235.17 |
| reject_neighbor_border_tg_as_gh | Neighbor TG point 0.02° east of GH | Reject | Reject / Reject | 422 | 211.98 |
| reject_outgroups_no_au_anchor | allow_outgroups lacking any AU isolate | Reject | Reject / Reject | 422 | 235.21 |
| reject_human_kmer_strict | Human 25-mer hit under strict_reject | Reject | Reject / Reject | 422 | 253.67 |
| reject_fastq_length_mismatch | Corrupted FASTQ seq vs. quality length | Reject | Reject / Reject | 422 | 259.29 |
| reject_missing_declared_file | Tier 3 parity: missing declared sequence file | Reject | Reject / Reject | 422 | 233.76 |

## B Structured-Filter and Access-Control Correctness (27 Queries)

Table 8 lists all 24 positive structured-filter/token checks and 3 negative zero-hit checks run at the 1,000-repository / 10,000-sample tier against SQL rules over visible (public and gated) records (with 10% private records excluded).

**Table 8:** All 27 structured-filter correctness queries on the 10,000-sample catalog (mean P@5 = 1.000, P@10 = 1.000, R@10 = 0.3157, 3/3 negative zero-hit checks, 0 private leaks).

| Query | Idx | Filter | SQL Oracle Rule | Rel. | Ret. | P@5 | P@10 | R@10 | MRR | NDCG |
|---|---|---|---|---|---|---|---|---|---|---|
| Nigeria | DS | country=NG | Vis. & country=NG | 25 | 20 | 1.0 | 1.0 | 0.400 | 1.0 | 1.0 |
| Malawi | DS | country=MW | Vis. & country=MW | 510 | 20 | 1.0 | 1.0 | 0.020 | 1.0 | 1.0 |
| Kenya | DS | country=KE | Vis. & country=KE | 101 | 20 | 1.0 | 1.0 | 0.099 | 1.0 | 1.0 |
| Uganda | DS | country=UG | Vis. & country=UG | 103 | 20 | 1.0 | 1.0 | 0.097 | 1.0 | 1.0 |
| Tanzania | DS | country=TZ | Vis. & country=TZ | 176 | 20 | 1.0 | 1.0 | 0.057 | 1.0 | 1.0 |
| Madagascar | DS | country=MG | Vis. & country=MG | 75 | 20 | 1.0 | 1.0 | 0.133 | 1.0 | 1.0 |
| Ghana | DS | country=GH | Vis. & country=GH | 18 | 18 | 1.0 | 1.0 | 0.556 | 1.0 | 1.0 |
| Senegal | DS | country=SN | Vis. & country=SN | 13 | 13 | 1.0 | 1.0 | 0.769 | 1.0 | 1.0 |
| Cameroon | DS | country=CM | Vis. & country=CM | 14 | 14 | 1.0 | 1.0 | 0.714 | 1.0 | 1.0 |
| Escherichia | DS | None | Visible *E. coli* cohorts | 900 | 20 | 1.0 | 1.0 | 0.011 | 1.0 | 1.0 |
| Ahmadu Bello | DS | None | Title/desc phrase match | 1 | 1 | 1.0 | 1.0 | 1.000 | 1.0 | 1.0 |
| Kilimanjaro | DS | None | Title/desc phrase match | 26 | 20 | 1.0 | 1.0 | 0.385 | 1.0 | 1.0 |
| Fondation Merieux | DS | None | Title/desc phrase match | 30 | 20 | 1.0 | 1.0 | 0.333 | 1.0 | 1.0 |
| DUBLIN | DS | None | Title/desc phrase match | 110 | 20 | 1.0 | 1.0 | 0.091 | 1.0 | 1.0 |
| Nigeria Escherichia | DS | country=NG | NG AND *E. coli* | 25 | 20 | 1.0 | 1.0 | 0.400 | 1.0 | 1.0 |
| Kenya Escherichia | DS | country=KE | KE AND *E. coli* | 101 | 20 | 1.0 | 1.0 | 0.099 | 1.0 | 1.0 |
| stool | Smp | None | source LIKE %stool% | 50 | 20 | 1.0 | 1.0 | 0.200 | 1.0 | 1.0 |
| blood | Smp | None | source LIKE %blood% | 101 | 20 | 1.0 | 1.0 | 0.099 | 1.0 | 1.0 |
| urine | Smp | None | source LIKE %urine% | 212 | 20 | 1.0 | 1.0 | 0.047 | 1.0 | 1.0 |
| feces | Smp | None | source LIKE %feces% | 242 | 20 | 1.0 | 1.0 | 0.041 | 1.0 | 1.0 |
| faeces | Smp | None | source LIKE %faeces% | 396 | 20 | 1.0 | 1.0 | 0.025 | 1.0 | 1.0 |
| Homo sapiens | Smp | None | host LIKE %homo sap.% | 8,684 | 20 | 1.0 | 1.0 | 0.001 | 1.0 | 1.0 |
| Bos taurus | Smp | None | host LIKE %bos tau.% | 3 | 3 | 1.0 | 1.0 | 1.000 | 1.0 | 1.0 |
| Lilongwe | Smp | None | admin1 LIKE %lilong.% | 2 | 2 | 1.0 | 1.0 | 1.000 | 1.0 | 1.0 |
| avian (Negative) | Smp | None | Zero-match isolation check | 0 | 0 | — | — | — | — | Pass |
| rodent (Negative) | Smp | None | Zero-match isolation check | 0 | 0 | — | — | — | — | Pass |
| NonExistentOutbreak99 | DS | None | Zero-match phrase check | 0 | 0 | — | — | — | — | Pass |

## C Complete Reference Sequence Accessions and Validation Inventory

Table 9 enumerates every NCBI RefSeq record, human sequencing benchmark, and ENA run identifier evaluated in this study.

**Table 9:** Complete NCBI RefSeq, human benchmark, and ENA sequencing accessions.

| Evaluation Group | Organisms / Targets | Exact Accession Identifiers |
|---|---|---|
| Bacterial RefSeq (7 seqs, 7 orgs) | *E. coli* K-12, *K. pneumoniae*, *M. tuberculosis*, *V. cholerae* chr1, *S. aureus*, *S. enterica*, *A. baumannii* | NC_000913, NC_016845, NC_000962, NC_002505, NC_007795, NC_003197, CP000521 |
| Parasite RefSeq (7 seqs, 4 orgs) | *P. falciparum* (chr1, apicoplast, mt, 18S rRNA), *T. brucei* chr1, *L. major* chr1, *S. mansoni* mt | NC_004325, NC_037282, NC_037526, XR_002966654, NC_026774, NC_001905, NC_002545 |
| Fungal RefSeq (1 seq, 1 org) | *Candida albicans* SC5314 chrR | NC_032096 |
| Viral RefSeq (7 seqs, 7 orgs) | Mpox, SARS-CoV-2, Lassa L, Zaire ebolavirus, RVFV L, Yellow fever, HIV-1 HXB2 | NC_063383, NC_045512, NC_004297, NC_002549, NC_014397, NC_002031, NC_001802 |
| Human Panel Construction | Human mtDNA, 18S rRNA, 28S rRNA, AluY (5,861 tiles) | NC_012920.1, NR_003286.4, NR_003287.4, U14574.1 |
| Independent Human Validation | *Homo sapiens* WGS run; single-copy *ACTB* mRNA | SRR30459529, NM_001101.5 |
| ENA African Screen | 5 clinical stool/WGS runs (SRR11015023 traced to *C. sabaeus*) | ERR17896308, SRR10973537, SRR11015023, SRR34287596, SRR39558023 |
| ENA Nigerian 613.83 MiB Storage | 3 paired-end runs (6 FASTQ.GZ files) + manifests | SRR9681899_1/2, SRR9822635_1/2, SRR9822643_1/2 |

## D Uniform 0.01% Seed Audit and Graded Free-Text BM25 Evaluation

Across the 30 uniform spike-in trials at *p* = 0.0001 (seeds `10100`–`10129`, 5 spiked reads each out of 50,000), 25/30 seeds had zero hits on both `head` and `multi_window`. Five trials placed at least one spike inside either the deterministic `head` range (0–1, 999) or the `multi_window` sample: trial 4 (`seed=10103`, positions [5277, 18859, 20714, 36158, 36164]; caught by `multi_window` reservoir at read `5277`), and trials 7, 8, 17, 28 (`seeds=10106, 10107, 10116, 10127`; single hits at reads `1307, 1375, 1652, 1668` caught by `head` outside the 0–999 prefix). An independent 200-seed check (`uniform_0001_convergence_200seed`) detected 38/200 (19.0%, CI [0.142, 0.250]).

Table 10 reports per-query results for Experiment 5b (free-text weighted BM25 without structured country filters).

**Table 10:** Experiment 5b free-text weighted BM25 evaluation across independent graded relevance labels (rel ∈ {0, 1, 2}; macro P@5 = 0.6945, R@10 = 0.9167, MRR = 1.000, graded NDCG@10 = 0.9200, 0 private leaks).

| **Free-Text Query** | **Returned Grades Order** | **P@5** | **R@10** | **MRR** | **NDCG** | **Ranking / Recall Observation** |
|---|---|---|---|---|---|---|
| malawi bloodstream | [2, 2, 1, 0] (priv. excluded) | 0.750 | 1.000 | 1.000 | 1.000 | Private holdout hidden; distractor last |
| ghana malaria resistance | [2, 2, 1, 0] | 0.750 | 1.000 | 1.000 | 1.000 | Title weight > benefit_sharing |
| kenya cholera outbreak | [2, 2, 1, 0] | 0.750 | 1.000 | 1.000 | 1.000 | Regional rel=1 beats SOP distractor |
| nigeria lassa hemorrhagic | [2, 2, 0, 1] | 0.750 | 1.000 | 1.000 | 0.987 | Ebola desc. distractor ranks above rel=1 |
| senegal pediatric diarrhea | [2, 0] (misses synonym) | 0.500 | 0.500 | 1.000 | 0.613 | Misses “childhood gastroenteritis” |
| south africa multidrug tuberculosis | [2, 0, 2] | 0.667 | 1.000 | 1.000 | 0.920 | Ethiopia desc. note interleaves at rank 2 |

